# Identification of stromal genes differentially expressed in lobular breast cancer highlights role for pregnancy-associated-plasma protein-A

**DOI:** 10.1101/2020.04.24.059386

**Authors:** Laura Gómez-Cuadrado, Hong Zhao, Margarita Souleimanova, Pernille Rimmer Noer, Arran K Turnbull, Claus Oxvig, Nicholas Bertos, J Michael Dixon, Morag Park, Andrew H Sims, Valerie G Brunton

## Abstract

**Background:** Invasive lobular carcinoma (ILC) is the second most common histological subtype of breast cancer and exhibits a number of clinico-pathological characteristics that are distinct from the more common invasive ductal carcinoma (IDC). Despite these differences, ILC is treated in the same way as IDC. We set out to identify alterations in the tumor microenvironment (TME) of ILC with potential clinical significance.

**Methods:** We used laser-capture microdissection (LCM) to separate tumor epithelium from stroma in 23 ER+ ILC samples. Gene expression analysis was used to identify genes that are enriched in the stroma of ILC, but not IDC or normal breast.

**Results:** 45 genes involved in regulation of the extracellular matrix (ECM) were enriched in the stroma of ILC, but not stroma from ER+ IDC or normal breast. Of these, 10 were expressed in cancer-associated fibroblasts (CAFs) and were increased in ILC compared to IDC in bulk gene expression datasets. *PAPPA* was the most enriched in the stroma compared to the tumor epithelial compartment in ILC. *PAPPA* encodes pregnancy-associated plasma protein-A (PAPP-A), a metalloproteinase that cleaves insulin-like binding protein-4 (IGFBP-4) increasing IGF-1 bioavailability and subsequent downstream signaling. Analysis of *PAPPA* and *IGF1* associated genes identified a paracrine signaling pathway and active PAPP-A was shown to be secreted from primary CAFs. Comprehensive survival analysis across 3000 breast cancers identified *PAPPA* as a potential ILC-specific prognostic marker.

**Conclusions:** This is the first study to demonstrate molecular differences in the TME between ILC and IDC and identifies PAPP-A, a CAF-derived proteinase, as a potential prognostic marker.

## Introduction

Invasive lobular breast cancer (ILC) accounts for around 5-15% of breast cancers and is the second most common histological subtype after invasive breast cancer of no specific type, commonly referred to as invasive ductal carcinoma (IDC). ILC is recognised to exhibit a number of clinico-pathological characteristics distinct from those of IDC [1, 2]. It has an increased propensity for multi-centricity, multi-focality and bilaterality, in addition to an unusual pattern of metastatic dissemination having a predilection for spread to the gastro-intestinal tract, peritoneum and ovary [3]. ILC is predominantly estrogen receptor (ER) and progesterone receptor positive, with low to absent expression of human epidermal growth factor receptor-2. Most patients with ILC are candidates for adjuvant endocrine treatment. Although response rates are initially good, an ILC diagnosis is associated with adverse long-term outcomes compared to IDC [4]. At the molecular level, ILC is defined by a loss or reduced expression of the cell-cell adhesion molecule E-cadherin, and several studies have further mapped the genomic landscape of ILC [5-8]. More recently, tumor-infiltrating lymphocyte populations have been profiled [9]. ILC is characterized by having a dense stroma which has a larger contact area with the tumor cells than in IDC, due in part to the difference in tumor growth pattern (“indian file” vs dense islands, respectively). However, little else is known about the composition of the stroma or the role of the surrounding tumor microenvironment (TME). The TME plays a critical role in tumor behaviour by influencing progression and metastatic spread, as well as therapeutic response [10, 11], and in breast cancer a stroma-derived prognostic predictor has been identified that stratifies disease outcome independently of clinical prognostic factors [12].

In this study we used laser-capture microdissection (LCM) to generate the first human ILC stromal gene set. A number of genes were more highly expressed in the stroma of ILC, but not that of IDC or normal breast. These genes included *PAPPA* encoding pregnancy-associated plasma protein-A (PAPP-A), a metalloproteinase that cleaves insulin-like binding protein-4 (IGFBP-4) when IGF-1 is bound to IGFBP-4 [13]. This results in a local increase in IGF-1 bioavailability and subsequent downstream signaling. Here we further show that active PAPP-A is secreted from cancer-associated fibroblasts (CAFs), and may represent an ILC-specific prognostic marker.

## Methods

All standard assays not detailed here are described in the Supplementary Methods (available online).

### Tissue processing for LCM and gene expression analysis

All samples were obtained from the McGill University Health Centre: Breast Cancer Functional Genomics Initiative Biobank, Montreal, Canada (Study identifiers SUR-99-780 and SUR-2000-966). Tissue collection, LCM, sample isolation, RNA extraction and microarray hybridization were carried out as previously described [14] and analyzed using the SurePrint G3 Human GE 8×60K microarray kit. Processed and raw data are available from Gene Expression Omnibus (GSE148398).

### Primary CAF dataset generation

All samples were obtained from the NHS Lothian Tissue Governance Committee, Edinburgh, United Kingdom, approval number 15/ES/0094. CAFs were isolated from five ILC and three IDC samples and total RNA extracted (Qiagen). RNA was biotinylated using the Illumina TotalPrep RNA Amplification kit (Ambion). Samples were run on Illumina HT-12 v4 BeadChips. Quantile normalisation was performed using the Lumi package. Processed and raw data are available from Gene Expression Omnibus (GSE148156).

### Statistical analysis

All statistical analyses were two-sided and p<0.05 was considered statistically significant. Differential gene expression analyses from LCM and CAF datasets were calculated using Rank Products in MeV [15]. Differences in gene expression in whole tissues were assessed by Wilcoxon test, in primary CAFs by Mann-Whitney-Wilcoxon test and in KEP tumor and CAFs by t-test. Mann-Whitney-Wilcoxon and fold change analysis were used to assess differences in gene expression between tumor and stroma in LCM datasets. RNAScope data were assessed by paired t-test. Correlation between *PAPPA* and *IGF1* was assessed by Pearson correlation and linear regression analysis. Comprehensive survival analysis was performed using the survival R package [16].

## Results

### Generation of an LCM-ILC dataset

LCM was performed on 23 ILC fresh frozen human samples. Three-quarters of tumors were Grade 2 (17, 74%), along with five Grade 1 and one Grade 3. RNA was isolated from tumor epithelium (TE) and tumor stroma (TS) compartments (TS defined here as primarily CAFs and matrix proteins, with the majority of immune cells being excluded). Gene expression data was generated for a total of 22 TE and 18 TS samples (Figure 1A; Supplementary Figure 1), including 17 matched TE and TS samples from the same patient. Two-class paired Rank Products analysis (percent false positive <0.01) identified 1,082 genes consistently more highly expressed in the TS and 837 in the TE when comparing TS versus TE. These genes clustered the samples by cell type (epithelium/stroma), showing successful microdissection of TE and TS compartments. REACTOME gene ontology annotation revealed up-regulation of genes involved in extracellular matrix (ECM) remodelling, collagen degradation and integrin cell surface interactions in TS compared to TE, while genes related to cell cycle, DNA replication and methylation were up-regulated in TE compared to TS compartments (Figure 1B).

**Figure 1:**
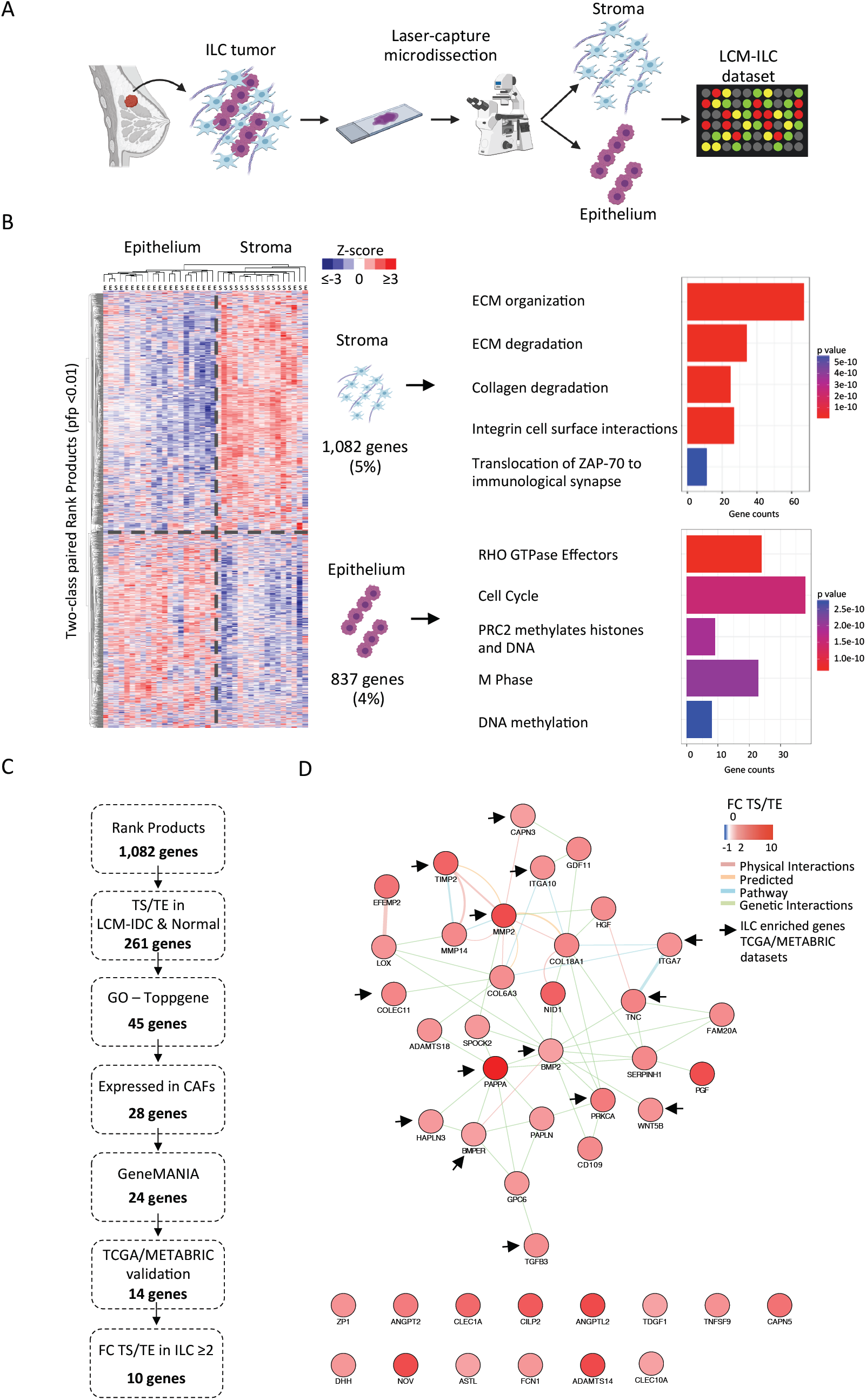
Identification of stromal genes enriched in ILC. A) Schematic representation of the LCM experimental design. B) Two-Class paired differential gene expression analysis and REACTOME pathway enrichment analysis of 22 tumor epithelial (TE) and 18 tumor stromal (TS) ILC samples dissected by LCM. Heatmap was generated using MultiExperiment Viewer (Rank Products, pfp < 0.01), Cluster3 and TreeView. Z-scored expression values. REACTOME bar plot was generated in R. C) Pipeline used for the selection of TS-ILC enriched genes from the LCM-ILC dataset. D) Interaction map of the 45 ECM associated genes from ToppGene was generated using GeneMANIA in Cytoscape. Colour represents the fold change (FC) expression in the TS compared to TE. Blue = FC TS/TE −1 – 0; White = FC TS/TE 0; Pink = FC TS/TE 1 – 2; Red = FC TS/TE 2 – 10. Network connectors represent physical interactions (pink), predicted (orange), pathway (blue) or genetic interactions (green). Arrows point to the 14 genes whose expression was significantly increased in ILC compared to IDC in TCGA and METABRIC datasets.

### Identification of TS-ILC enriched genes

An analysis pipeline was set up to identify genes up-regulated in the TS compared to TE in ILC, but not IDC or normal breast (Figure 1C). First, the list of 1,082 genes differentially expressed in our TS LCM-ILC dataset was applied to previously reported LCM-IDC (GSE68744) [17] and LCM-normal (GSE4823) [14] datasets. This identified 261 genes that were only increased in the TS compared to TE in ILC. ToppGene gene ontology analysis (https://toppgene.cchmc.org/) identified 45 of these genes to be involved in significant pathways (Benjamini-Hochberg adjusted p-value <0.05), all related to the ECM (Figure 1D; Supplementary Figure 2). Network analysis revealed 30 interconnected genes, including matrix proteins (*COL6A3, COL18A1, TNC, EFEMP2*), proteoglycans (*SPOCK2, PAPLN, HAPLN3, GPC6*), proteinases and their regulators (*MMP2, TIMP2, MMP14, CAPN3, ADAMTS18, SERPINH1, PAPPA*) and integrin subunits (*ITGA7* and *ITGA10*). A number of growth factors, including those of the TGFβ superfamily (*PGF, GFD11, HGF, TGFB3, BMP2*), were also identified. The analysis highlighted physical interactions of *MMP2* with *TIMP2, MMP14, COL6A3, COL18A1* and *CAPN3*, all involved in ECM organization. In addition, *BMP2* and *PAPPA* were the two main hubs of genetic interactions [18] (Figure 1D). The expression of these 45 stromal genes was examined in ILC and IDC ER+ samples from the METABRIC [19] and The Cancer Genome Atlas (TCGA; http://cancergenome.nih.gov/) bulk mRNA datasets (Table 1). *PAPPA, PRKCA, TGFB3, ITGA10, ITGA7, CLEC1A, CLEC10A* and *PAPLN* were up-regulated in ILC compared to IDC in all 3 datasets (METABRIC, TCGA RNA-Seq and TCGA microarray), highlighting the importance of these genes in the stroma of lobular carcinoma (Supplementary Figure 3).

**Table 1.** List of the 45 ECM genes from the ToppGene GO analysis in ILC, IDC- and normal-LCM datasets and whole tissue datasets. Fold changes for LCM datasets are TS/TE. Bulk datasets indicate significant gene expression difference between subtypes, high is indicated, Mann-Whitney-U *p<0.05, ***p<0.0001.

|  | Genes | LCM datasets – tissue fold change |  |  | Whole tissue datasets – subtype significant |  |  |
| --- | --- | --- | --- | --- | --- | --- | --- |
|  |  | ILC<br>(GSE148398) | IDC<br>(GSE68744) | NORMAL<br>(GSE4823) | METABRIC | TCGA<br>microarray | TCGA RNA-Seq |
| Genes in<br>CAF<br>dataset<br>(28) | PAPPA | 9.14 | 1.05 | 0.01 | ILC* | ILC* | ILC* |
|  | ANGPTL2 | 6.29 | 1.26 | 0.50 | ILC* | - | ILC*** |
|  | NOV | 6.21 | 0.93 | 0.03 | - | ILC*** | ILC*** |
|  | MMP2 | 6.20 | 1.01 | -0.14 | - | ILC* | ILC*** |
|  | NID1 | 4.90 | 1.21 | 0.20 | - | - | - |
|  | TIMP2 | 4.77 | 0.97 | 0.12 | - | - | ILC* |
|  | CAPN5 | 4.02 | 1.12 | 0.18 | ILC* | - | ILC*** |
|  | EFEMP2 | 3.88 | 0.76* | 0.00 | IDC*** | - | ILC*** |
|  | PRKCA | 3.52 | 1.16 | 0.36 | ILC* | ILC* | ILC*** |
|  | TNC | 3.13 | 1.07 | -0.20 | - | ILC* | ILC* |
|  | COL18A1 | 2.95 | 0.98 | 0.05 | IDC* | - | ILC*** |
|  | COLEC11 | 2.95 | 1.02 | 0.64 | ILC* | - | - |
|  | MMP14 | 2.81 | 1.02 | -0.01 | - | - | - |
|  | SERPINH1 | 2.78 | 0.88 | -0.17 | IDC* | - | - |
|  | CD109 | 2.77 | 1.52 | 0.44 | IDC* | - | - |
|  | GDF11 | 2.67 | 1.03 | -0.01 | IDC*** | - | - |
|  | TGFB3 | 2.59 | 1.60 | 0.00 | ILC*** | ILC* | ILC*** |
|  | COL6A3 | 2.53 | 0.98 | -0.03 | - | - | - |
|  | TNFSF9 | 2.36 | 1.04 | 0.08 | IDC* | - | - |
|  | ITGA10 | 2.35 | 1.14 | 0.10 | ILC*** | ILC*** | ILC*** |
|  | ITGA7 | 2.33 | 0.90 | 0.34 | ILC* | ILC*** | ILC*** |
|  | WNT5B | 2.30 | 1.23 | 0.00 | - | - | ILC* |
|  | LOX | 2.28 | 1.25 | 0.07 | - | - | - |
|  | GPC6 | 2.04 | 1.00 | 0.16 | IDC* | - | IDC*** |
|  | HAPLN3 | 1.98 | 0.99 | -0.06 | - | - | ILC*** |
|  | CAPN3 | 1.89 | 1.01 | 0.00 | - | - | ILC*** |
|  | BMP2 | 1.86 | 1.14 | -0.05 | - | ILC* | ILC*** |
|  | BMPER | 1.86 | 1.06 | -0.04 | - | - | ILC*** |
| Genes<br>not in<br>CAF<br>dataset<br>(17) | ADAMTS14 | 6.37 | 1.18 | 0.03 | IDC* | - | - |
|  | PGF | 6.08 | 0.96 | -0.06 | - | - | ILC*** |
|  | CILP2 | 5.35 | 1.03 | -0.01 | - | - | ILC*** |
|  | CLEC1A | 4.51 | 1.06 | 0.06 | ILC*** | ILC* | ILC*** |
|  | ANGPT2 | 3.46 | 1.00 | 0.07 | IDC* | - | IDC* |
|  | FAM20A | 2.66 | 1.01 | 0.10 | - | ILC* | ILC*** |
|  | HGF | 2.58 | 1.05 | 0.20 | - | ILC* | ILC*** |
|  | ZP1 | 2.35 | 1.05 | 0.03 | - | - | - |
|  | ADAMTS18 | 2.28 | 1.08 | -0.01 | - | - | ILC*** |
|  | FCN1 | 2.07 | 0.76* | 0.48 | - | - | ILC*** |
|  | MUC19 | 2.04 | 0.98 | -0.01 | - | - | - |
|  | DHH | 2.04 | 0.89 | -0.19 | - | - | ILC*** |
|  | SPOCK2 | 2.02 | 1.14 | 0.11 | - | ILC* | ILC*** |
|  | PAPLN | 1.98 | 1.15 | 0.21 | ILC*** | ILC* | ILC*** |
|  | CLEC10A | 1.84 | 0.92* | 0.43 | ILC* | ILC* | ILC*** |
|  | ASTL | 1.82 | 1.10 | -0.01 | - | IDC* | ILC* |
|  | TDGF1 | 1.81 | 1.31 | -0.27 | - | - | ILC*** |

### ILC-specific expression of primary cancer-associated fibroblasts

In order to perform functional studies, gene expression profiling of primary CAFs from ILC and IDC tumors was performed (Figure 2A and B; Supplementary Figure 4). Rank Products analysis (percent false positive <0.05) identified 1,027 genes differentially expressed between ILC- and IDC-derived CAFs; approximately half (485) were up-regulated in ILC CAFs compared to IDC CAFs (Figure 2C) and were enriched for ECM-associated genes such as *COL6A3* and *FN1*, along with genes involved in glycolysis and focal adhesion, as well as members of the TGFβ signaling pathway (Figure 2D).

**Figure 2:**
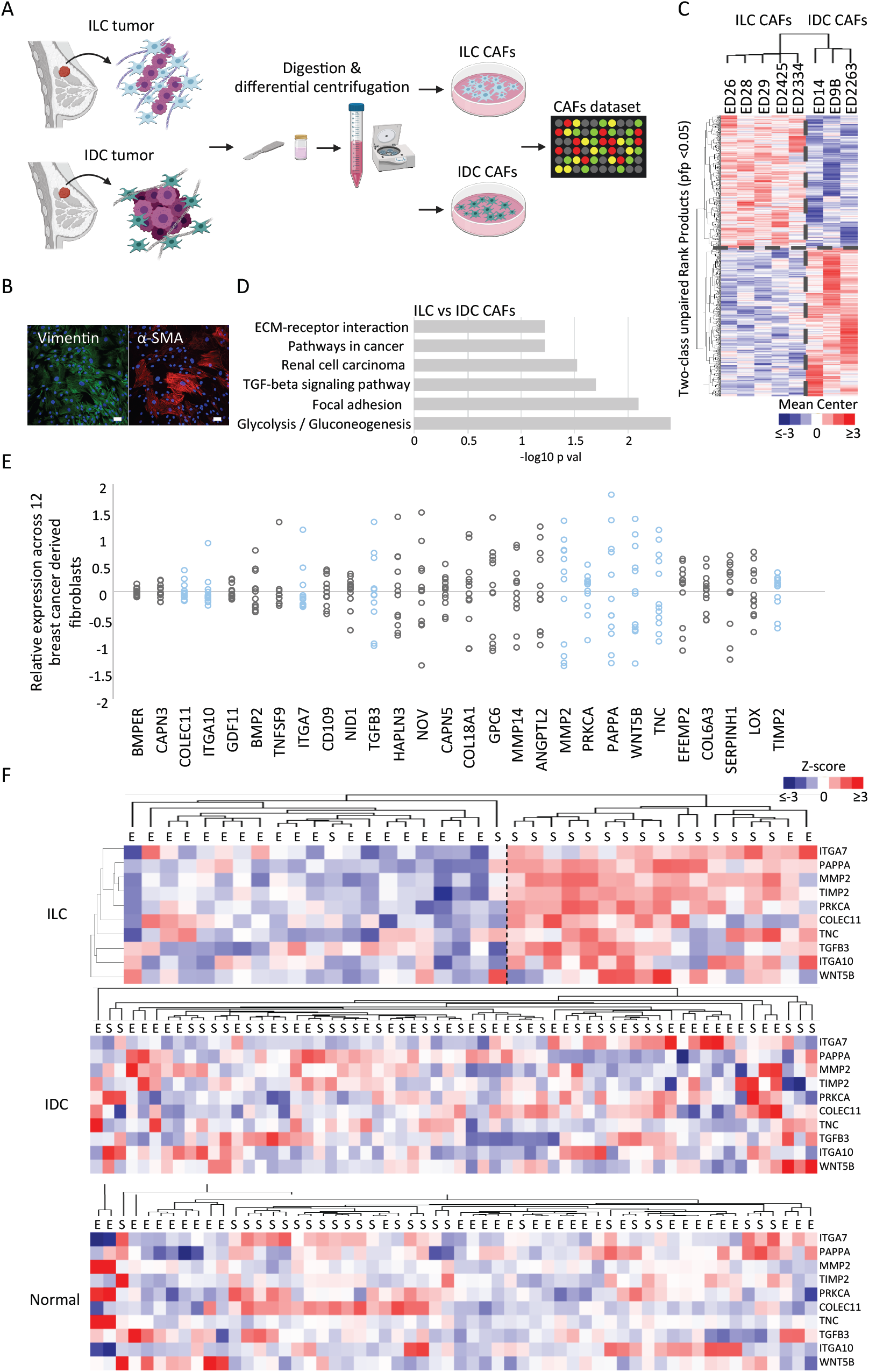
Generation of a primary CAF dataset. A) Schematic representation of primary CAFs dataset generation. B) Primary ED2425 ILC CAFs stained for the mesenchymal markers vimentin and α-SMA. Scale bar = 50μm. C) Two-Class unpaired differential gene expression analysis. Heatmap was generated using MultiExperiment Viewer (Rank products, pfp < 0.05), Cluster3 and TreeView. Mean-centred expression values. D) Differentially expressed genes and pathways between ILC and IDC CAFs was performed using DAVID Bioinformatics Resources (KEGG pathways). E) Relative expression of 28 genes expressed in primary CAFs. Marked in blue the 10 ILC CAF-associated genes. F) Heatmaps of CAF-associated genes in LCM-ILC, -IDC, - Normal datasets were generated from Z-scored values using Cluster3 and TreeView. Tissue samples are indicated as E=epithelium, S=stroma.

Of the 45 ILC-specific stromal genes identified by LCM, 28 were expressed in the CAF dataset (Table 1; Figure 2E). GeneMANIA (http://genemania.org/) analysis identified that 24 of these 28 genes were in the same pathway, or have/predicted genetic or physical interactions (Supplementary Figure 5). The majority (14/24) of these genes were significantly up-regulated in ILC compared to IDC (p<0.05) in at least one of the published bulk datasets (Table 1). Clustering the 10 genes with a TS/TE fold change >2 clearly showed increased expression in the TS of ILC, but not in IDC or normal breast (Figure 2F).

### *PAPPA* is predominantly expressed in the stroma of ILC

*PAPPA* was taken forward for further analysis as it showed the greatest fold change expression in the stromal compared to epithelial compartments in ILC (FC TS/TE >9) (Table 1). *PAPPA* encodes PAPP-A, a secreted metalloproteinase that cleaves IGFBP-4, releasing bioactive IGF-1 that can initiate downstream signaling via the IGF-1R [20]. PAPP-A activity can be inhibited by non-covalent or covalent complex formation with endogenous inhibitors stanniocalcin-1 (STC1) or −2 (STC2), respectively [21, 22] (Figure 3A). To verify that *PAPPA* was expressed predominantly by CAFs and not by tumor cells, we analyzed *PAPPA* transcripts by RNAScope. Results confirmed higher levels in CAFs compared to tumor cells, although epithelial *PAPPA* transcripts were also seen in some of the tumors (Figure 3B). We then examined the expression of *PAPPA* and functionally related genes in the LCM-ILC, -IDC and -normal datasets (Figure 3C). In ILC both *PAPPA* and *IGF1* were significantly more highly expressed in the stroma compared to the epithelium (p ≤ 0.0001), while *IGF1R* was found predominantly in the tumor epithelium (p ≤ 0.05), suggesting the presence of a potential paracrine activation loop. In IDC and normal breast tissue, *IGF1* was also predominantly expressed in the stroma (p ≤ 0.0001 and p ≤ 0.01 respectively), whereas *PAPPA* was expressed in both stromal and epithelial compartments (Figure 3C). Interestingly, we found a clear positive correlation between *PAPPA* and *IGF1* in ILC (r = 0.64, p<0.0001) which was not observed in IDC and normal tissue (Figure 3D). As few cell lines that represent ILC are available, we first examined *PAPPA* expression across three integrated breast cancer cell line datasets [23]. *PAPPA* was low or undetectable in all luminal cell lines, including two reported ILC lines SUM44-PE and MDA-MB-134VI (Figure 4A). qPCR confirmed that *PAPPA* was not expressed in the SUM44-PE and MDA-MB-134VI ILC lines or the T47D and MCF-7 ER+ IDC lines. Analysis of 11 primary patient-derived ILC CAFs, 5 IDC CAFs, and HCI 013, an ILC patient-derived xenograft showed that *PAPPA* and *IGF1* were expressed exclusively in CAFs (Figure 4B; Supplementary Figure 6). In contrast, *IGF1R* was mostly expressed by tumor cells (Figure 4B). We also separated tumor epithelial cells from CAFs in tumors derived from a mouse model of ILC driven by loss of *Trp53* and *Cdh1* (Supplementary Figure 7A) [24]. qPCR results showed that *Pappa, Igf1* and *Stc1* were only expressed in the CAFs, while *Igf1r, Stc2* and *Igfbp4* were expressed in both tumor cells and CAFs (Supplementary Figure 7B). This supports a wider paracrine signaling role for PAPP-A in luminal tumors.

**Figure 3:**
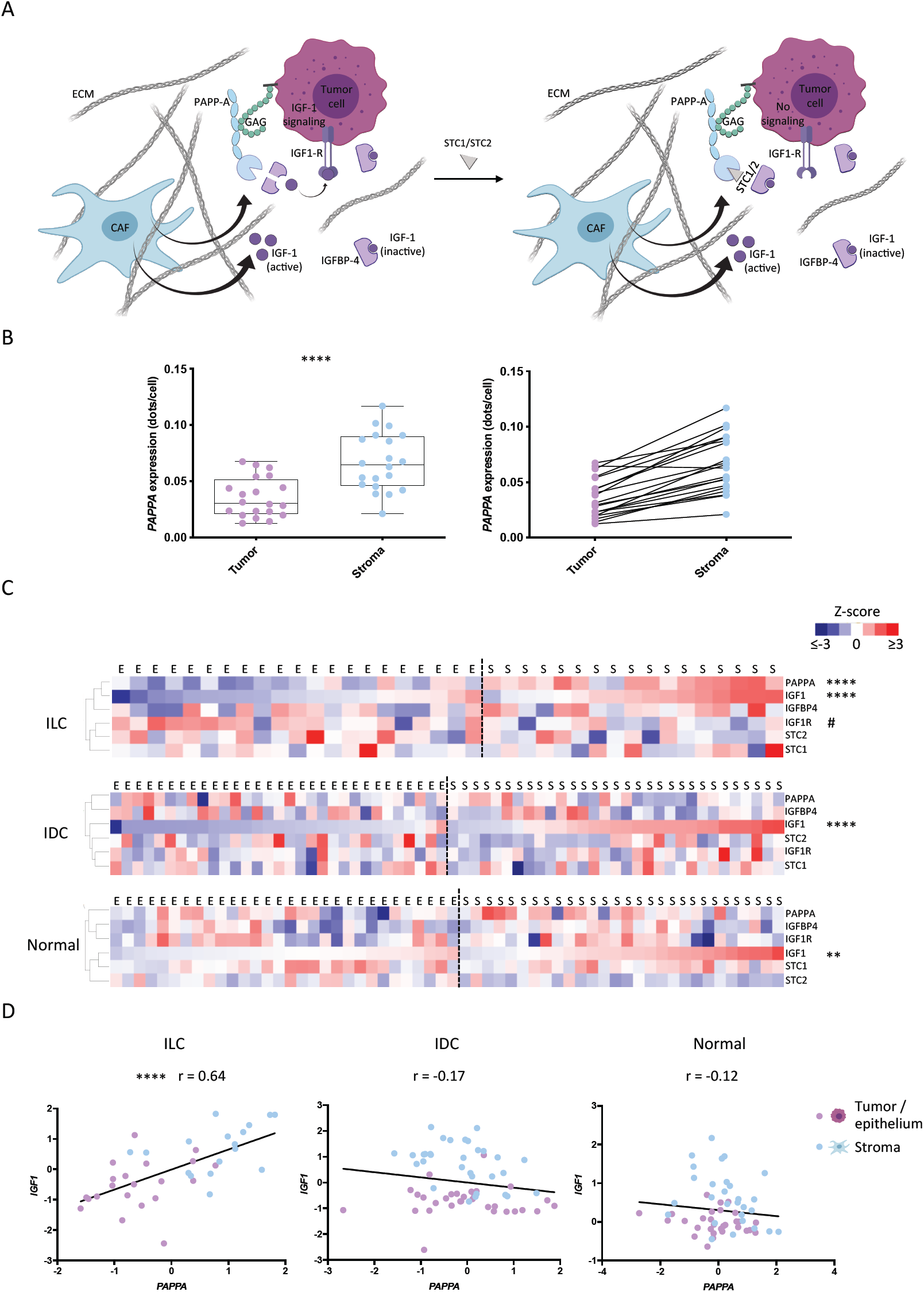
PAPPA is a stromal-derived factor in ILC. A) Schematic representation of PAPP-A function in lobular breast cancer. B) RNAScope was performed in 20 of the primary samples used for generation of the LCM-ILC dataset. Samples were analyzed using QuPath. Left - Boxplots representing expression of PAPPA (dots per cell) in the tumor and stroma. Paired t-test **** p value <0.0001. Right - Trend lines show an increase in PAPPA in the stroma of ILC samples compared to the tumor cells. C) Heatmaps representing PAPPA and its regulators in the epithelial (E) and stromal (S) compartments of ILC-, IDC- (GSE68744) and normal- (GSE4823) LCM datasets. Heatmaps were generated from Z-scored values using Cluster3 and TreeView. Adjusted p values were calculated using Wilcox test in R. ** p ≤ 0.01, **** p ≤ 0.0001, # p ≤ 0.05. * Means up in stroma vs epithelium, # means up in the epithelium vs stroma. D) Scatterplots representing PAPPA – IGF1 correlation in ILC-, IDC-(GSE68744) and normal- (GSE4823) LCM datasets. Pearson correlation and linear regression analyses were performed in GraphPad. **** p ≤ 0.0001.

**Figure 4:**
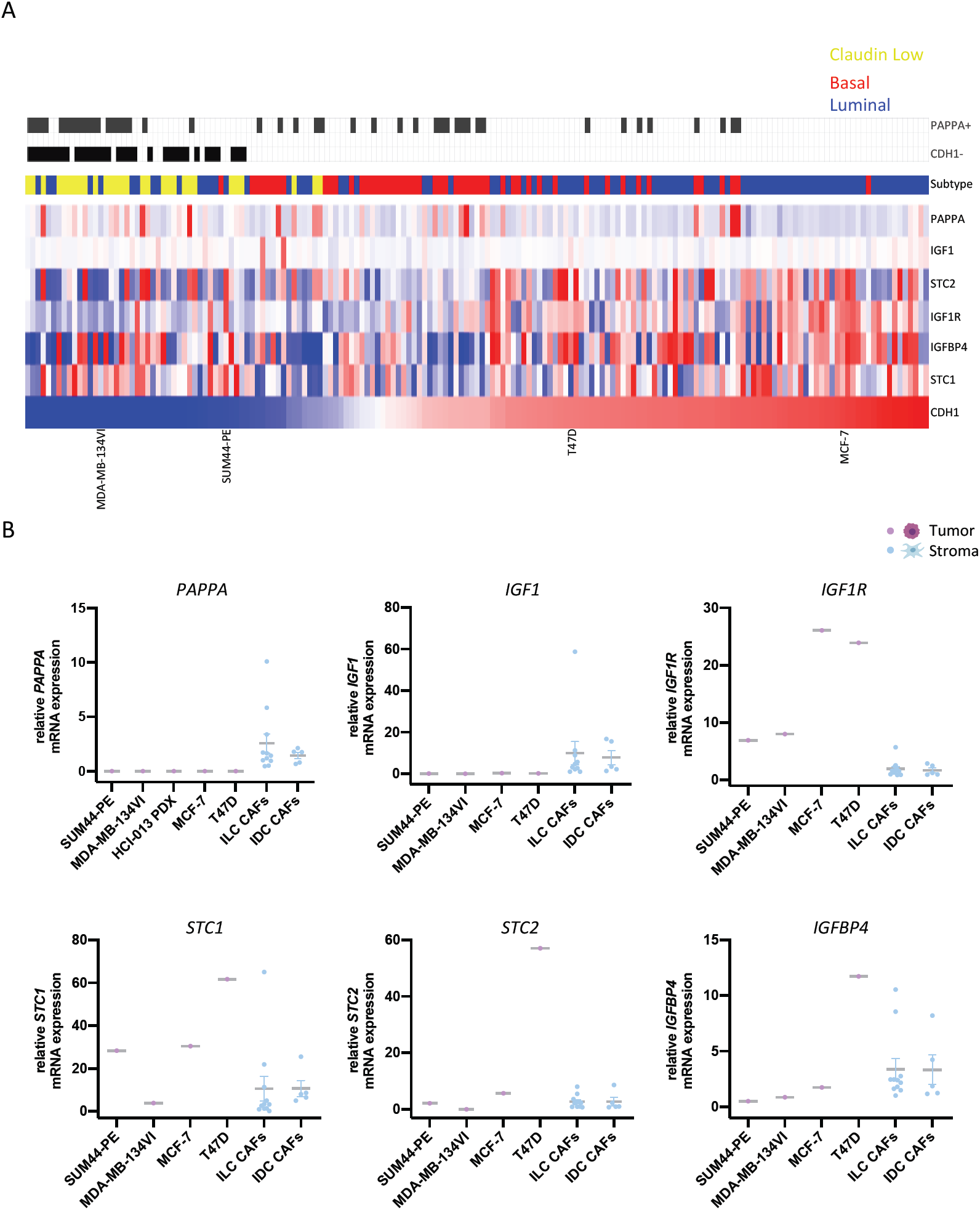
PAPPA in in vitro models of ILC. A) Heatmap representing the expression of PAPPA and associated genes across three integrated panels of breast cancer cell lines following batch correction, ranked by CDH1 expression. Blue=Luminal, Red=Basal, Yellow=Claudin-low. Grey bars indicate samples where the detection call for PAPPA is assigned as ‘present’ and those tumors where CDH1 is ‘absent’. B) Expression of PAPPA, IGF1, IGF1R, SCT1, SCT2 and IGFBP4 by qPCR in ILC & IDC human cell lines and primary CAFs. Each sample was analyzed at 3 different passage numbers, and its average represented as the relative mRNA expression to ED30 primary ILC CAFs. Line represents the mean with SEM.

### PAPP-A secreted by CAFs is active

To establish whether PAPP-A was secreted from CAFs, we analyzed conditioned media (CM) and confirmed that PAPP-A was secreted by the CAFs but not by the tumor cells (Figure 5A). PAPP-A needs to be active in order to cleave IGFBP-4 and liberate IGF-1. CM from the CAFs was able to cleave recombinant IGFBP-4, indicating that non-complexed active PAPP-A was present in the media (Figure 5B). To confirm that the IGFBP-4 fragments generated by the CAF CM were a result of PAPP-A activity, the CM was treated with a PAPP-A inhibitory antibody, (mAb 1/41) [25]. Pre-incubation with mAb 1/41 reduced levels of the cleaved IGFBP-4 fragment to those in the control lane, showing that the observed cleavage of IGFBP-4 is due to PAPP-A present in the CM (Figure 5C).

**Figure 5:**
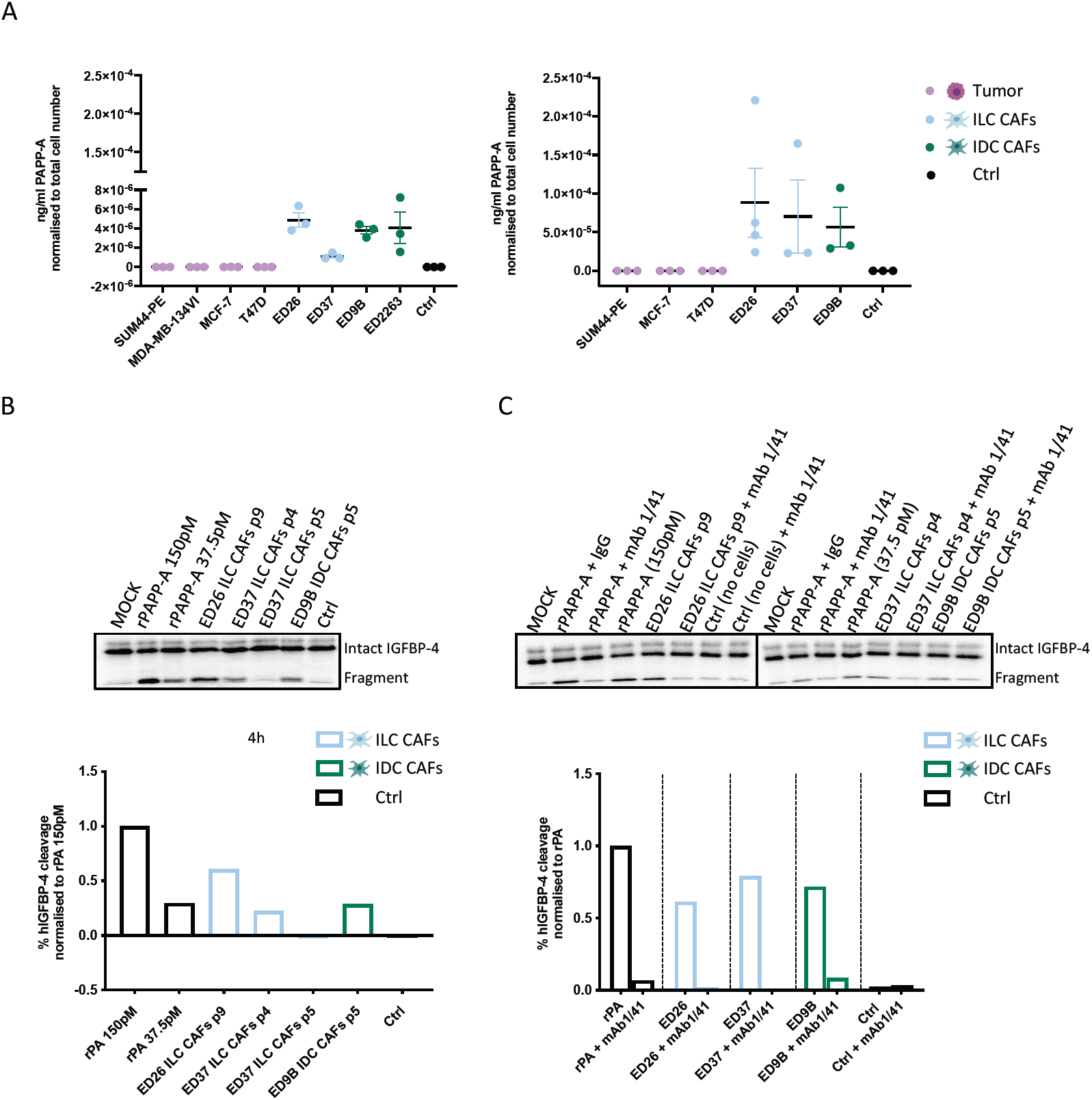
Active PAPP-A is secreted from CAFs. A) ELISA of PAPP-A secreted from a series of ILC and IDC cell lines and primary CAFs. Conditioned media (CM) was collected after 24 hours (left panels) or after 48 hours from the tumor cells and 96 hours from the CAFs (right panels). PAPP-A concentration was normalized to total cell number, n = 3. B) PAPP-A mediated IGFBP-4 cleavage. CM from ILC and IDC primary CAFs was incubated with radiolabeled IGFBP-4 for 1, 2 and 4 hours. Recombinant PAPP-A (rPAPP-A) was used as a positive control. Quantification at 4 hours is represented as the percentage of human IGFBP-4 cleavage normalised to recombinant PAPP-A (150 pM). C) IGFBP-4 cleavage was measured in the presence of PAPP-A inhibitory antibody (mAb 1/41) or isotype control (IgG) in CM from ILC and IDC CAFs. Quantification at 4 hours is represented as the percentage of human IGFBP-4 cleavage normalised to the corresponding recombinant PAPP-A (150 pM or 37 pM). p denotes passage number of the CAFs.

### *PAPPA* expression is positively correlated with *IGF1* and negatively with *IGF1R* and is elevated in *CDH1-*, claudin-low tumors

Investigation of the expression of *PAPPA* and related genes in large cohorts of breast cancers confirmed positive correlations between *PAPPA* and *IGF1* and negative correlations with *IGF1R* and *CDH1* (Figure 6A and 6B). An integrated compendium of 17 Affymetrix datasets representing 2999 breast cancers [23] revealed that *PAPPA* is detectably expressed in 36% of tumors, the highest proportion of which were of the claudin-low subtype, which also had significantly higher levels of *PAPPA* expression (Figure 6A). Somewhat surprisingly, 7% of ILCs in the METABRIC dataset are classified as claudin-low, compared to only 4% of IDCs and *PAPPA* expression was significantly higher in tumors where *CDH1* was undetectable (Figure 6B).

**Figure 6:**
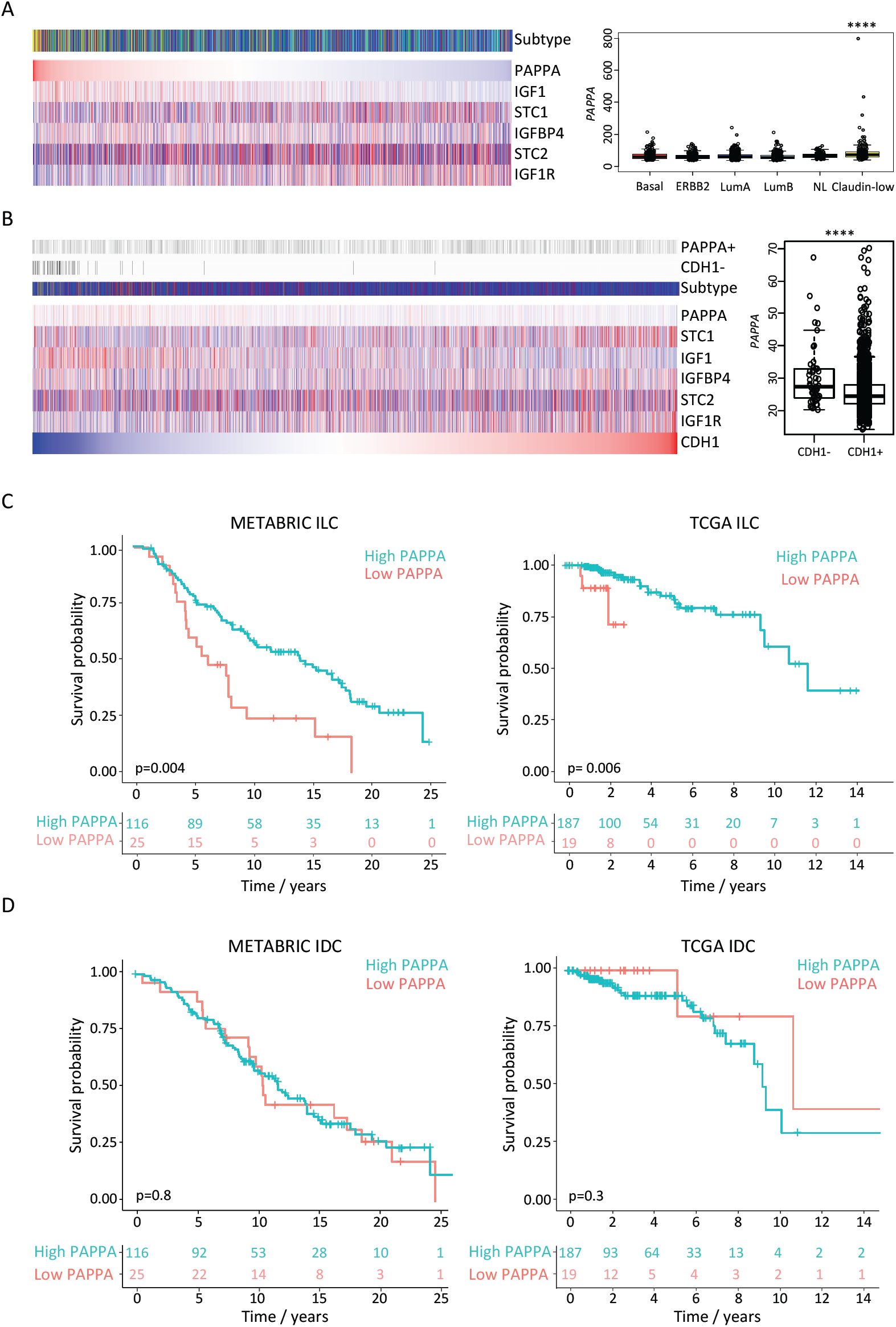
Low *PAPPA* expression in lobular, but not ductal breast cancers is associated with poor outcomes. A) Breast tumors of the METABRIC cohort ranked by PAPPA expression indicates positive correlation with IGF1, negative correlation with IGF1R and highest expression in claudin-low subtype tumors. **** p ≤ 0.0001. Heatmap shows log2 mean-centred expression values, Red=high, blue=low. Subtypes are indicated by the coloured bar above the heatmap. red=basal, purple=ERBB2, dark blue=Luminal A, light blue=luminal B, green=normal-like, yellow=claudin-low. B) Ranked expression of 2999 breast tumors by CDH1 expression indicates that PAPPA is significantly higher in CDH1-tumors. **** p ≤ 0.0001. Grey bars above the heatmap indicate detection calls for tumors where PAPPA is called ‘present’ and CDH1 is called absent. C) Kaplan Meier plots for overall survival showing the most significant cut-points for PAPPA expression in ILC tumors from the METABRIC and TCGA studies. D) Representative Kaplan Meier plots for 142 and 206 randomly sampled ER+ IDCs from METABRIC and TCGA to compare with the same number of ILCs shown in panel C.

### Low *PAPPA* expression in ILC, but not IDC is associated with poor outcomes

To examine whether *PAPPA* relates to prognosis of breast cancers, we performed comprehensive survival analysis using the survivALL R package [16]. Low *PAPPA* was significantly (p<0.05) associated with worse overall survival for a large proportion (807/1903) of all possible (n-1) cut-points for 1904 breast cancers in the METABRIC cohort. However, none of the cut-points for *PAPPA* expression were significantly associated with overall survival across the 1098 breast cancers of the TCGA dataset (Supplementary Figure 8A). There are a number of major differences between these cohorts, not least the length of follow-up and the date of diagnosis. Restricting analysis to the 142 and 206 ILCs of the METABRIC and TCGA cohorts respectively identified a number of cut-points that were significantly associated with overall survival in both cohorts (Figure 6C; Supplementary Figure 8B). To compare this with IDCs from the two cohorts, analysis was limited to the same numbers (142 and 206) of ER+ IDC tumors with 10-fold random sampling, but on no occasion were any significant cut-points identified (Figure 6D; Supplementary Figure 8C). Taken together, these results suggest that reduced or absent *PAPPA* is an ILC-specific prognostic marker.

## Discussion

In this study we provide the first comprehensive analysis of the TME in ILC. This identified a set of 45 genes enriched in the stroma of ILC but not in the stroma of IDC or normal breast, that were associated with ECM regulation. CAFs are an important component of the TME that regulate ECM deposition and secrete cytokines and growth factors to control metastatic spread and therapeutic response [26]. However, little is known about the involvement of CAFs in the biology of ILC, although one study has reported an increased CAF density in ILC compared to matched IDC samples [27], while differences in collagen deposition and alignment have also been reported in ILC [28, 29]. Our analysis of primary CAFs from ILC and IDC shows differences in a number of genes involved in ECM interactions and signaling pathways indicating that CAFs are able to differentially influence the TME in ILC compared to IDC. A wider analysis of CAFs from ILC will be required to fully understand their importance and also whether the heterogeneity and different sub-populations of CAFs that exist in other breast cancers is pertinent to ILC [30, 31]. Interestingly, four of the stromal-derived ECM associated genes (*PRKCA, ITGA10, NOV, WNT5B*) that we identified in ILC were found in the *reactive-like* ILC subtype described by Ciriello and colleagues [5]; these *reactive-like* ILC tumors largely associate with the *reactive* subgroup identified by TCGA that are characterized by strong microenvironment and CAF signaling [32].

We have focused here on *PAPPA*, which encodes an IGF-promoting proteinase. Further analysis of *PAPPA* and functionally associated genes showed that both *PAPPA* and *IGF1* were predominantly expressed in the stroma of ILC while *IGF1R* was expressed within the tumor epithelium. Together with the demonstration that active PAPP-A is secreted from CAFs strongly supports the existence of a paracrine signaling pathway in ILC. Further analysis of a panel of breast cancer cell lines showed that *PAPPA* was low or undetectable in all luminal cell lines in support of a wider PAPP-A paracrine rather than autocrine signaling axis in other breast cancer subtypes. Of note, *PAPPA* was elevated in the *reactive* ILC breast cancer subtype (characterised by proteins produced by the TME and CAFs) when analyzing both TCGA and METABRIC datasets (Supplementary Figure 8D). However, of the few immunohistochemical studies reporting on PAPP-A in breast tumors, only expression in the tumor epithelium is recorded. A recent study demonstrated greater extent and intensity of PAPP-A expression in luminal B than luminal A tumors, although sample numbers were low [33]. PAPP-A has also been proposed to be a tumor suppressor following the discovery that it is epigenetically silenced in breast cancer precursor lesions [34]. Comprehensive survival analysis in this study of the two largest publicly available breast cancer gene expression datasets indicate that reduced *PAPPA* is associated with worse outcomes, but this finding appears to be limited to lobular, rather than ductal, breast cancers. Interestingly, *PAPPA* levels were negatively associated with *CDH1*, and *PAPPA* was significantly higher in tumors where *CDH1* was undetected, while the highest proportion of *PAPPA* positive tumors were of the claudin-low subtype, which are characterized by low or absent expression of *CDH1* [35], suggesting a potential functional link between PAPP-A and E-cadherin.

Previously we have shown that loss of E-cadherin promotes hypersensitization of PI3K/Akt pathway activation in response to IGF1, independent of PAPP-A [36], as well as oncogenic mutations in the PI3K/Akt pathway that are prevalent in ILC [5]. This is consistent with other reports demonstrating that E-cadherin-mediated adhesion negatively regulates IGF1R activation [37, 38]. Furthermore, in breast cancer models, loss of E-cadherin and the subsequent activation of IGF1R signaling results in increased sensitivity to dual IGF1R/Insulin receptor inhibitors, and Akt inhibitors that target downstream receptor pathway activation, even in the presence of activating *PIK3CA* mutations [36, 37]. Interestingly, increased expression of *IGF1* is seen in ILC compared to IDC [27, 36, 39], consistent with reported pathway activation [36, 37]. Together, these data suggest that patients with ILC may benefit from treatments targeting the IGF1 signaling pathway. Whether this would be more effective when used in combination with inhibitors of PI3K and Akt in the context of activating mutations in the PI3K/Akt pathway remains to be established.

Although a number of strategies to target the IGF1/IGF1R signaling axis have been tested, results in the clinical setting have been disappointing with a number of contributory factors including effects on systemic glucose metabolism and associated metabolic toxicities and lack of predictive biomarkers [40]. Indirect targeting of IGF1R signaling via inhibition of PAPP-A proteolytic activity may provide a viable alternative by reducing the levels of bioactive IGF1 specifically in the local microenvironment of tumors that express high levels of PAPP-A. Consistent with this are reports that treatment with a monoclonal antibody that blocks the proteolytic activity of PAPP-A reduces tumor growth in both lung cancer and Ewing sarcoma models, which express high levels of PAPP-A within the tumor cells [25, 41]. In addition, in a series of patient derived ovarian tumor xenografts, antitumor activity of the PAPP-A inhibitory antibody correlated with PAPP-A expression [42]. Interestingly, in the 4T1 murine model of breast cancer, which expresses E-cadherin, expression of a protease-resistant IGFBP-4, or treatment with recombinant PAPP-A resistant IGFBP-4, inhibited tumor growth and metastasis via inhibition of PAPP-A secreted by the surrounding stromal tissue [43, 44]. A recent study found that circulating levels of PAPP-A in patients with breast cancer were independently prognostic for recurrence-free and overall survival [45]. Thus, circulating levels of PAPP-A may provide a route to patient stratification when considering targeting PAPP-A.

Overall this study demonstrates that PAPP-A is an important stromal factor in ILC. Intriguingly, the correlation between E-cadherin and PAPP-A together with the reported role for E-cadherin in regulating IGF1R activation indicates that there is a wider role for PAPP-A in regulating breast cancer growth.

## Notes

The authors have no conflicts of interest to disclose.

### Author contributions

Initiated the project: LGC, MP, AHS, VGB. Sample preparation and collection: LGC, HZ, MS, JMD. Performed experiments and interpreted data: LGC, PRN, CO. Statistics and bioinformatics: LGC, AHS, AKT, NB. Supervised research: MP, AHS, VGB. Wrote the manuscript: LGC, AHS, VGB. Approved the manuscript: all authors.

The KEP tumors were a kind gift from Seth Coffelt and were generated by Jos Jonkers and Karen de Visser. We thank Frances Rae, Lorna Renshaw and Jane Keys for assistance in collection of primary tissue.

## References

1. Iorfida M, Maiorano E, Orvieto E, et al. Invasive lobular breast cancer: subtypes and outcome. Breast Cancer Res Treat 2012;133(2):713–23.

2. Thomas M, Kelly ED, Abraham J, et al. Invasive lobular breast cancer: A review of pathogenesis, diagnosis, management, and future directions of early stage disease. Semin Oncol 2019;46(2):121–132.

3. Ferlicot S, Vincent-Salomon A, Medioni J, et al. Wide metastatic spreading in infiltrating lobular carcinoma of the breast. Eur J Cancer 2004;40(3):336–41.

4. Pestalozzi BC, Zahrieh D, Mallon E, et al. Distinct clinical and prognostic features of infiltrating lobular carcinoma of the breast: combined results of 15 International Breast Cancer Study Group clinical trials. J Clin Oncol 2008;26(18):3006–14.

5. Ciriello G, Gatza ML, Beck AH, et al. Comprehensive Molecular Portraits of Invasive Lobular Breast Cancer. Cell 2015;163(2):506–19.

6. Desmedt C, Zoppoli G, Gundem G, et al. Genomic Characterization of Primary Invasive Lobular Breast Cancer. J Clin Oncol 2016;34(16):1872–81.

7. Michaut M, Chin SF, Majewski I, et al. Integration of genomic, transcriptomic and proteomic data identifies two biologically distinct subtypes of invasive lobular breast cancer. Sci Rep 2016;6:18517.

8. Shah SP, Morin RD, Khattra J, et al. Mutational evolution in a lobular breast tumour profiled at single nucleotide resolution. Nature 2009;461(7265):809–13.

9. Desmedt C, Salgado R, Fornili M, et al. Immune Infiltration in Invasive Lobular Breast Cancer. J Natl Cancer Inst 2018;110(7):768–776.

10. Junttila MR, de Sauvage FJ. Influence of tumour micro-environment heterogeneity on therapeutic response. Nature 2013;501(7467):346–54.

11. Quail DF, Joyce JA. Microenvironmental regulation of tumor progression and metastasis. Nat Med 2013;19(11):1423–37.

12. Finak G, Bertos N, Pepin F, et al. Stromal gene expression predicts clinical outcome in breast cancer. Nat Med 2008;14(5):518–27.

13. Conover CA, Oxvig C. PAPP-A and cancer. J Mol Endocrinol 2018;61(1):T1–t10.

14. Finak G, Sadekova S, Pepin F, et al. Gene expression signatures of morphologically normal breast tissue identify basal-like tumors. Breast Cancer Res 2006;8(5):R58.

15. Wang YE, Kutnetsov L, Partensky A, et al. WebMeV: A Cloud Platform for Analyzing and Visualizing Cancer Genomic Data. Cancer Res 2017;77(21):e11–e14.

16. Pearce DA, Nirmal AJ, Freeman TC, et al. Continuous Biomarker Assessment by Exhaustive Survival Analysis. bioRxiv 2018;https://doi.org/10.1101/208660.

17. Oh EY, Christensen SM, Ghanta S, et al. Extensive rewiring of epithelial-stromal co-expression networks in breast cancer. Genome Biol 2015;16:128.

18. Lin A, Wang RT, Ahn S, et al. A genome-wide map of human genetic interactions inferred from radiation hybrid genotypes. Genome Res 2010;20(8):1122–32.

19. Curtis C, Shah SP, Chin SF, et al. The genomic and transcriptomic architecture of 2,000 breast tumours reveals novel subgroups. Nature 2012;486(7403):346–52.

20. Oxvig C. The role of PAPP-A in the IGF system: location, location, location. J Cell Commun Signal 2015;9(2):177–87.

21. Jepsen MR, Kloverpris S, Mikkelsen JH, et al. Stanniocalcin-2 inhibits mammalian growth by proteolytic inhibition of the insulin-like growth factor axis. J Biol Chem 2015;290(6):3430–9.

22. Kloverpris S, Mikkelsen JH, Pedersen JH, et al. Stanniocalcin-1 Potently Inhibits the Proteolytic Activity of the Metalloproteinase Pregnancy-associated Plasma Protein-A. J Biol Chem 2015;290(36):21915–24.

23. Moleirinho S, Chang N, Sims AH, et al. KIBRA exhibits MST-independent functional regulation of the Hippo signaling pathway in mammals. Oncogene 2013;32(14):1821–30.

24. Derksen PW, Liu X, Saridin F, et al. Somatic inactivation of E-cadherin and p53 in mice leads to metastatic lobular mammary carcinoma through induction of anoikis resistance and angiogenesis. Cancer Cell 2006;10(5):437–49.

25. Mikkelsen JH, Resch ZT, Kalra B, et al. Indirect targeting of IGF receptor signaling in vivo by substrate-selective inhibition of PAPP-A proteolytic activity. Oncotarget 2014;5(4):1014–25.

26. Houthuijzen JM, Jonkers J. Cancer-associated fibroblasts as key regulators of the breast cancer tumor microenvironment. Cancer Metastasis Rev 2018;37(4):577–597.

27. Nakagawa S, Miki Y, Miyashita M, et al. Tumor microenvironment in invasive lobular carcinoma: possible therapeutic targets. Breast Cancer Res Treat 2016;155(1):65–75.

28. Burke K, Tang P, Brown E. Second harmonic generation reveals matrix alterations during breast tumor progression. J Biomed Opt 2013;18(3):31106.

29. Natal RA, Paiva GR, Pelegati VB, et al. Exploring Collagen Parameters in Pure Special Types of Invasive Breast Cancer. Sci Rep 2019;9(1):7715.

30. Bartoschek M, Oskolkov N, Bocci M, et al. Spatially and functionally distinct subclasses of breast cancer-associated fibroblasts revealed by single cell RNA sequencing. Nat Commun 2018;9(1):5150.

31. Costa A, Kieffer Y, Scholer-Dahirel A, et al. Fibroblast Heterogeneity and Immunosuppressive Environment in Human Breast Cancer. Cancer Cell 2018;33(3):463-479.e10.

32. Network CGA. Comprehensive molecular portraits of human breast tumours. Nature 2012;490(7418):61–70.

33. Mansfield AS, Visscher DW, Hart SN, et al. Pregnancy-associated plasma protein-A expression in human breast cancer. Growth Horm IGF Res 2014;24(6):264–7.

34. Loddo M, Andryszkiewicz J, Rodriguez-Acebes S, et al. Pregnancy-associated plasma protein A regulates mitosis and is epigenetically silenced in breast cancer. J Pathol 2014;233(4):344–56.

35. Prat A, Parker JS, Karginova O, et al. Phenotypic and molecular characterization of the claudin-low intrinsic subtype of breast cancer. Breast Cancer Res 2010;12(5):R68.

36. Teo K, Gomez-Cuadrado L, Tenhagen M, et al. E-cadherin loss induces targetable autocrine activation of growth factor signalling in lobular breast cancer. Sci Rep 2018;8(1):15454.

37. Nagle AM, Levine KM, Tasdemir N, et al. Loss of E-cadherin Enhances IGF1-IGF1R Pathway Activation and Sensitizes Breast Cancers to Anti-IGF1R/InsR Inhibitors. Clin Cancer Res 2018;24(20):5165–5177.

38. Qian X, Karpova T, Sheppard AM, et al. E-cadherin-mediated adhesion inhibits ligand-dependent activation of diverse receptor tyrosine kinases. Embo j 2004;23(8):1739–48.

39. Bertucci F, Orsetti B, Negre V, et al. Lobular and ductal carcinomas of the breast have distinct genomic and expression profiles. Oncogene 2008;27(40):5359–72.

40. Ekyalongo RC, Yee D. Revisiting the IGF-1R as a breast cancer target. NPJ Precis Oncol 2017;1.

41. Heitzeneder S, Sotillo E, Shern JF, et al. Pregnancy-Associated Plasma Protein-A (PAPP-A) in Ewing Sarcoma: Role in Tumor Growth and Immune Evasion. J Natl Cancer Inst 2019;111(9):970–982.

42. Becker MA, Haluska P, Jr., Bale LK, et al. A novel neutralizing antibody targeting pregnancy-associated plasma protein-a inhibits ovarian cancer growth and ascites accumulation in patient mouse tumorgrafts. Mol Cancer Ther 2015;14(4):973–81.

43. Ryan AJ, Napoletano S, Fitzpatrick PA, et al. Expression of a protease-resistant insulin-like growth factor-binding protein-4 inhibits tumour growth in a murine model of breast cancer. Br J Cancer 2009;101(2):278–86.

44. Smith YE, Toomey S, Napoletano S, et al. Recombinant PAPP-A resistant insulin-like growth factor binding protein 4 (dBP4) inhibits angiogenesis and metastasis in a murine model of breast cancer. BMC Cancer 2018;18(1):1016.

45. Espelund U, Renehan AG, Cold S, et al. Prognostic relevance and performance characteristics of serum IGFBP-2 and PAPP-A in women with breast cancer: a long-term Danish cohort study. Cancer Med 2018;7(6):2391–2404.

