## Supplementary methods and figures for "Identification of stromal genes differentially expressed in lobular breast cancer highlights role for pregnancy-associated-plasma protein-A"

**Tissue processing for LCM and gene expression analysis**

Tissues were obtained from patients who had previously provided written, informed consent for sample collection to a Research Ethics Board- (REB) approved biobanking initiative (MUHC study identifiers SUR-99-780 and SUR-2000-966). 23 fresh frozen samples were micro-dissected into epithelium and stroma using a PixCell IIe LCM system (Arcturus) and RNA was then isolated (PicoPure RNA isolation kit, Arcturus) and concentrated (LabConco). RNA was amplified using the T-7 based RiboAmp HS Plus RNA amplification kit (Arcturus) and labelled with Cy3 dye (Turbo labeling kit Cy3, Life Technologies) for two-color microarray-based gene expression hybridization and scanning (Agilent). Raw data was normalized, log2-transformed, filtered, and quality control analysis carried out. HGNC gene annotations were applied and the mean was taken when multiple probes were present for the same gene. The batch effect was corrected using ComBat [1]. Gene expression of TS differentially expressed genes from our ILC-LCM dataset were applied to previously reported LCM-IDC (GSE68744) [2] and LCM-normal (GSE4823) [3] datasets, which comprise 32 IDC ER+ paired TE-TS samples and 58 normal breast paired epithelium-stroma samples respectively.

**RNAScope**

FFPE sections were obtained from the 23 ILC samples used for the generation of the LCM-ILC dataset. Each section was stained with a positive control probe Hs-PPIB to check the quality of the RNA. 20 out of 23 samples had good RNA integrity and were stained for human *PAPPA* (NM_002571.5) targeting 1274-2258. Expression of *PAPPA* was quantified in QuPath version 0.2.0. 6 x 1 $mm^2$ regions of interest were analyzed per section and the number of dots per cell in the tumor and stroma calculated.

**Cell lines and generation of conditioned media**

SUM44-PE (BioIVT), MDA-MB-134VI, MCF-7, T47D (all ATCC), KEP epithelial cells and fibroblasts (provided by Seth Coffelt, Cancer Research UK Beatson Institute, Glasgow) and primary human CAFs (isolated from tumors obtained via the NHS Lothian Tissue Governance Committee under approval 15/ES/0094) were grown at 37 ºC in humidified air incubators with 5% $CO_2$. All CAFs were seeded on collagen-coated plates using 1:30 PureCol (Advanced BioMatrix). Cells were authenticated via STR profiling at Culture Collections, UK using the Promega Powerplex 16 HS kit and confirmed to be mycoplasma-negative by qPCR performed at the MRC Human Genetics Unit, The University of Edinburgh. To collect conditioned media (CM) cells were serum starved and the media collected after 24 and 48 hours (tumor cells) and 96 hours (CAFs). The CM was centrifuged at 4 ºC at 1600 rpm for 10 min to remove cell debris and filtered using a 0.22 μm filter.

**PAPP-A ELISA**

Conditioned media from SUM44-PE, MDA-MB-134VI, MCF-7, T47D, and primary ILC and IDC CAFs was used to detect levels of secreted PAPP-A in the media using a highly sensitive PAPP-A picoELISA kit as previously described [4]. PAPP-A concentration was normalized to total cell number. Data analyzed in GraphPad. n = 3.

**IGFBP-4 cleavage assay**

Conditioned media from ILC and IDC primary CAFs was tested for PAPP-A ability to cleave IGFBP-4 as previously described [5]. Briefly, samples were incubated with radiolabeled ($^{125}$I) IGFBP-4 for 1, 2 and 4 hours. Cleavage products were separated by 10-20% SDS-PAGE and visualized by autoradiography. The degree of cleavage was quantified using a Typhoon imaging system (GE Healthcare), and background levels were subtracted. In some reactions, purified mAb1/41 antibody was added in known amounts as specified.

**Western blotting**

Detection of PAPP-A and STC2 in the conditioned media of primary ILC and IDC CAFs was assessed. 3-8% Novex Tris-Acetate gels were used for the separation of the samples in NuPAGE Tris-Acetate SDS running buffer at 120V for 1 h 20 min. PA*S2, a mixture of recombinant PAPP-A dimer (~400 kDa) and recombinant PAPP-A-STC2 complex (~500 kDa) was used as a control. The gel was transferred to a PVDF membrane by a wet transfer at 30V for 1h or 500mA for 1h 30min in a Tris-glycine buffer (pH = 8) with 20% EtOH. Membrane was blocked in freshly made 2% Tween20 in $dH_2O$ for 20 min, washed in TBS-T x3. Primary antibody (1:2000) was incubated in TBS-T with 0.5% foetal bovine serum overnight at room temperature Secondary antibody (anti-rabbit 1:2000) was incubated for 60 min in TBS-T with 0.5% foetal bovine serum at room temperature. Membranes were developed with ECL using the BioRad imaging system.

**qRT-PCR**

RNA was isolated from ILC and IDC cell lines and primary CAFs as well as from tumors isolated from the HCI-013 PDX model (a gift from Dr Alana Welm, Utah) using the RNeasy kit (Qiagen). RNA was converted into cDNA using the SuperScript First-Strand kit (Invitrogen). qPCR was performed with a SYBR Select Mastermix (Life Technologies) and the ΔΔCt was calculated as the relative expression to ED30 CAFs or KEP CAFs. n = 3. Graphs were generated in GraphPad and the differences in gene expression were assessed by Mann-Whitney Wilcoxon test and t-test. Primer sequences and melting temperatures (Tm) are summarised in Supplementary Table 1.

#### Immunofluorescence

Cells were grown on glass cover-slips and fixed with 4% paraformaldehyde for 10 min at room temperature. Fixed samples were stained for vimentin (1:200, 5741S, Cell Signaling Technologies), α-SMA (1:200, M0851, Dako), cytokeratin18 (1:200, 4548S, Cell Signaling Technologies), cytokeratin14 (1:200, in-house), estrogen receptor (1:200, ab16660, Abcam), PR (1:100, M3569, Dako), E-cadherin (1:100, 3195S, Cell Signaling Technologies), P-cadherin (1:150, AF761, R&D) overnight at 4 ºC. Alexa Fluor 488 and 549 (Invitrogen) secondary antibodies were applied for 45 min at room temperature. Cover-slips were attached to slides using Vectashield with DAPI (Vector Laboratories). Fluorescent images were captured using Olympus FV1000 confocal microscope and Fluoview software and processed using ImageJ (National Institutes of Health).

#### Illustrations

Illustrations were created with BioRender.com

#### Primers used for RT-qPCR analysis

| Primers | Forward 5' - 3' | Reverse 5' - 3' | Species | Tm |
|---|---|---|---|---|
| **hGAPDH** | GGACCTGACCTGCCGTCTAG | TGGTGCTCAGTGTAGCCCAG | human | Tm = 56-62 ºC |
| **hIGF1** | CTTCAGTTCGTGTGTGGAGACAG | CGCCCTCCGACTGCTG | human | Tm = 59 ºC |
| **hIGF1R** | CTCCTGTTTCTCTCCGCCG | ATAGTCGTTGCGGATGTCGAT | human | Tm = 57 ºC |
| **hIGFBP4** | CCTCTACATCATCCCCATCC | GGTCCACACACCAGCACTT | human | Tm = 56 ºC |
| **hPAPPA** | AGCCAGCAGCATCCCAGGTGT | CGCCCGGAGCCAAAAAGTGGT | human | Tm = 62 ºC |
| **hSTC1** | ATTCCCACCAACAAAATCCA | CCGTTCTAAAGGGATCCACA | human | Tm = 56 ºC |
| **hSTC2** | ACTACTCAACTCTGCCGTCC | ACGCTTGGTTTCTTGGTGTC | human | Tm = 56 ºC |
| **mGAPDH** | CCACTCACGGCAAATTCAACGGCA | ACCAGTAGACTCCACGACATACTC | mouse | Tm = 56 ºC |
| **mIGF1** | CTGGTGGATGCTCTTCAGTTCG | TGCTTTTGTAGGCTTCAGTGGG | mouse | Tm = 58 ºC |
| **mIGF1R** | GGAGAAGCCCATGTGTGAG | GTCGTGGATAACGAAGCCATC | mouse | Tm = 57 ºC |
| **mIGFBP4** | GGAAAGGAATGGGGTGAGGA | AATATGGGGACGGAGGCAAA | mouse | Tm = 56 ºC |
| **mPAPPA** | CACTTGGGCGGTATTGTCTT | TGGGTTGGTATCATTGCAGA | mouse | Tm = 56 ºC |
| **mSTC1** | CCCAATCACTTCTCCAACAGA | GAAGAGGCTGGCCATGTTG | mouse | Tm = 56 ºC |
| **mSTC2** | GACCCTCTGGAAGCAGTGAG | ACACATCCAGCGTGTGACAT | mouse | Tm = 60 ºC |

### Supplementary Figures

#### Supplementary Figure 1

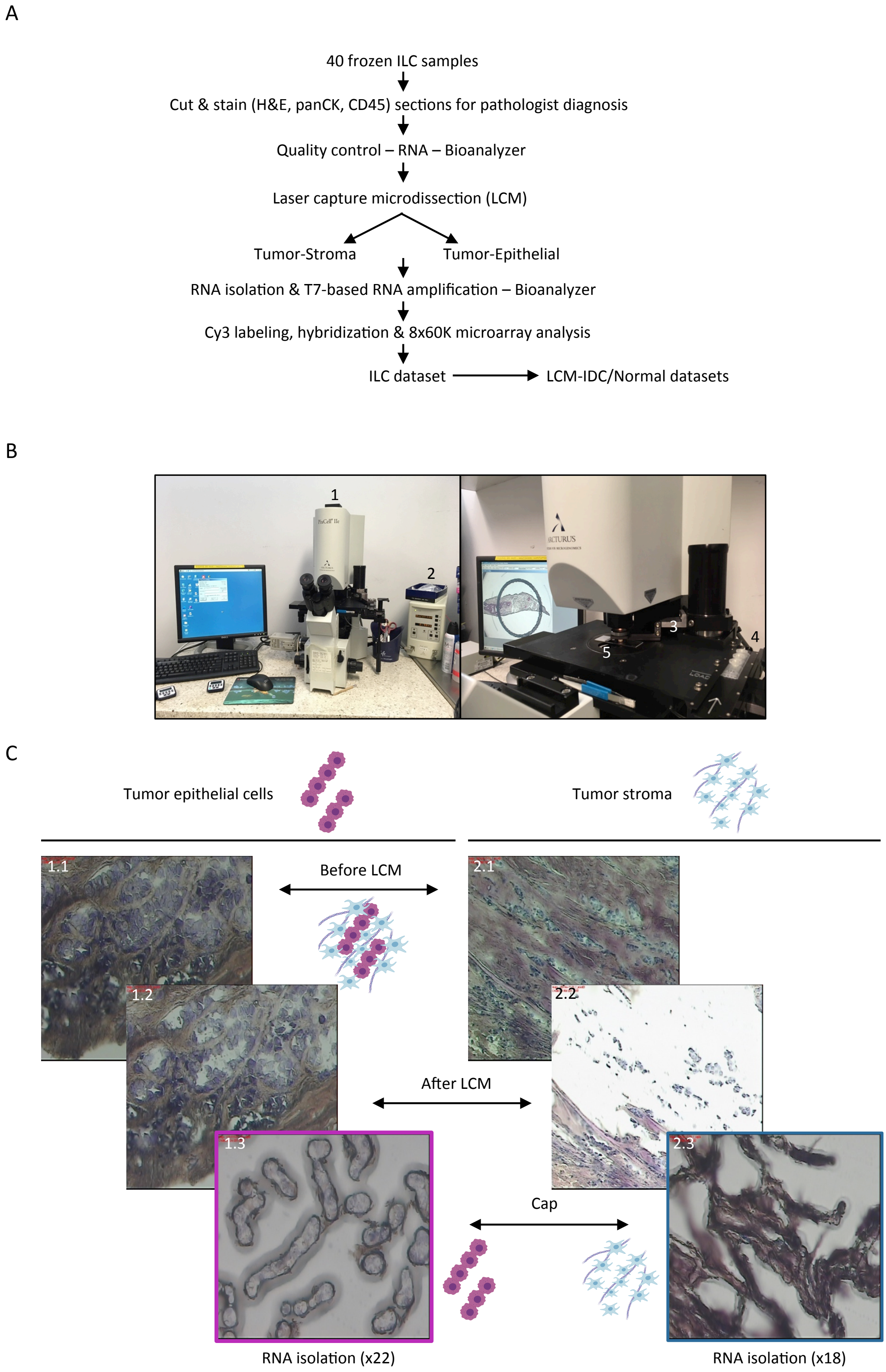


**Supplementary Figure 1: Overview of laser-capture microdissection technique.** A) Workflow used for the generation of a lobular gene expression dataset comprised of ILC epithelial (TE) and stromal cells (TS). B) PixCell IIe Laser Capture Microdissection microscope (1). An infrared laser (2) is used to lift the cells of interest. A handle (3) is used to place the HS Capture cap (4) on the top of the HistoBond stained slide (5). C) Images of two ILC tissue samples being processed. Left: isolation of tumor epithelial cells. Right: isolation of tumor stromal cells. 1.1 & 2.1) Tissue before LCM; 1.2 & 2.2) Tissue after LCM; 1.3 & 2.3) Caps after LCM. A total of 22 and 18 epithelial and stromal samples were isolated respectively from 23 ILC samples, 17 of them paired-matched.

#### Supplementary Figure 2

| | Pathway | Source | q-value FDR B&H | 45 genes merged |
|---|---|---|---|---|
| Toppgene | Ensemble of genes encoding extracellular matrix and extracellular matrix-associated proteins | MSigDB C2 BIOCARTA (v6.0) | 7.13E-05 | ADAMTS14, ADAMTS18, ANGPT2, ANGPTL2, ASTL, BMP2, BMPER, CAPN3, CAPN5, CD109, CILP2, CLEC10A, CLEC1A, COL18A1, COL6A3, COLEC11, DHH, EFEMP2, FAM20A, FCN1, GDF11, GPC6, HAPLN3, HGF, ITGA10, ITGA7, LOX, MMP14, MMP2, MUC19, NID1, NOV, PAPLN, PAPPA, PGF, PRKCA, SERPINH1, SPOCK2, TDGF1, TGFB3, TIMP2, TNC, TNFSF9, WNT5B, ZP1 |
| | Extracellular matrix organization | BioSystems: REACTOME | 1.32E-04 | |
| | Ensemble of genes encoding ECM-associated proteins including ECM-affilaited proteins, ECM regulators and secreted factors | MSigDB C2 BIOCARTA (v6.0) | 6.98E-03 | |

**Supplementary Figure 2: Gene ontology analysis of stromal genes enriched in ILC.** ToppGene gene ontology analysis of 261 genes (Benjamini-Hochberg adjusted p value <0.05) identified 45 genes involved in significant pathways related to the ECM.

### Supplementary Figure 3

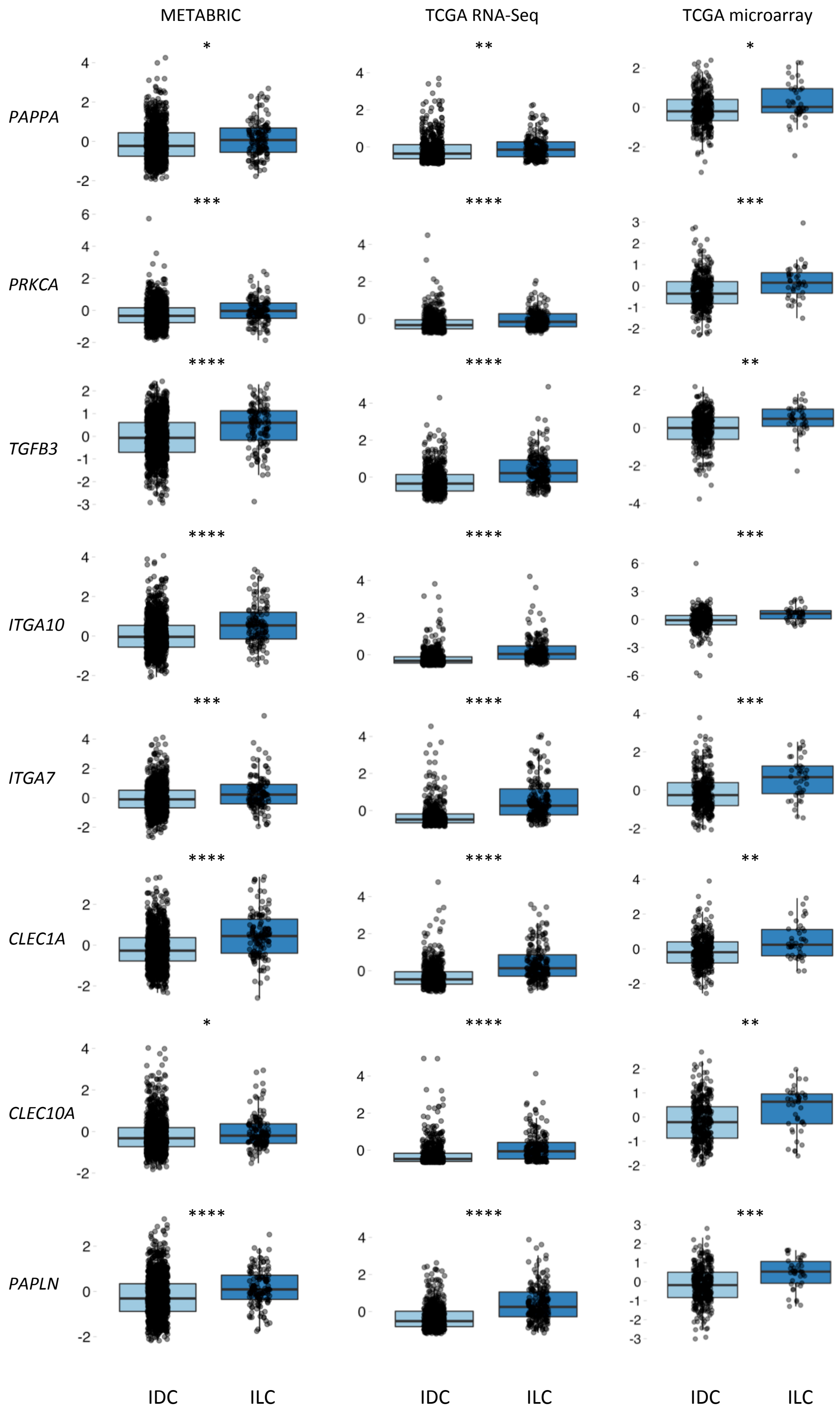

**Supplementary Figure 3: Expression of stromal genes enriched in ILC analysed in METABRIC, TCGA RNA-Seq and TCGA microarray datasets.** Boxplots of expression for TS-ILC enriched genes from METABRIC (Illumina v3 microarray) and TCGA provisional (RNA-Seq v2 and TCGA microarray) mRNA expression datasets (expression Z-scores; Z-score threshold ±2). Analysis was reduced to ILC and IDC ER-positive samples only. Boxplots display the median (line), 25th and 75th percentiles (box) and 1.5 × interquartile range (whiskers). Light blue, IDC; dark blue, ILC. * $p \leq 0.05$, ** $p \leq 0.01$, *** $p \leq 0.001$, **** $p \leq 0.0001$; Wilcoxon test. Boxplots were generated using the ggplot2 package in R.

#### Supplementary Figure 4

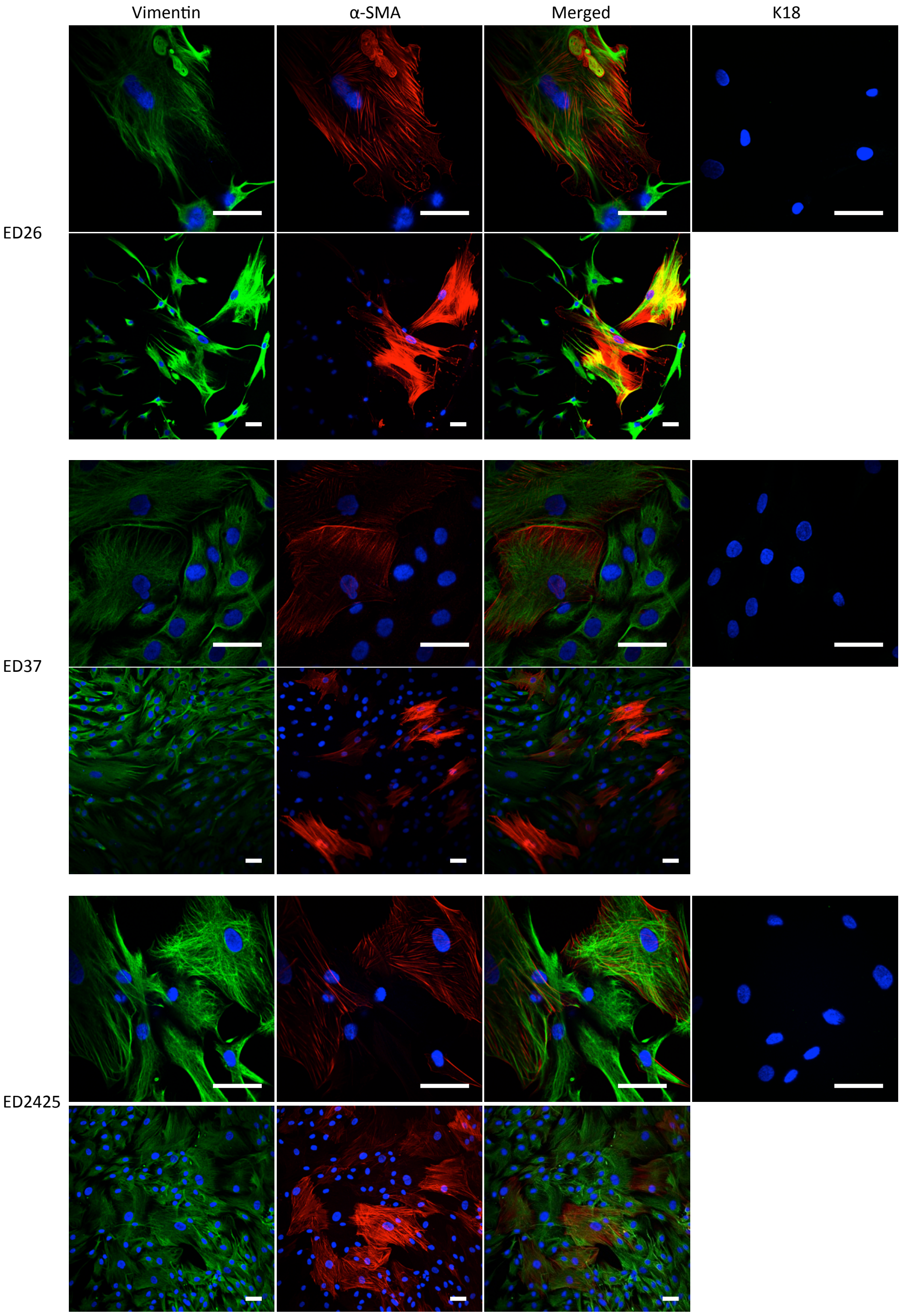


**Supplementary Figure 4: Characterization of primary CAFs.** Characterization of primary CAFs stained for the mesenchymal markers vimentin and α-SMA, and the epithelial marker keratin18. Examples provided for three ILC derived CAF lines. Scale bar = 50 μm.

Supplementary Figure 5

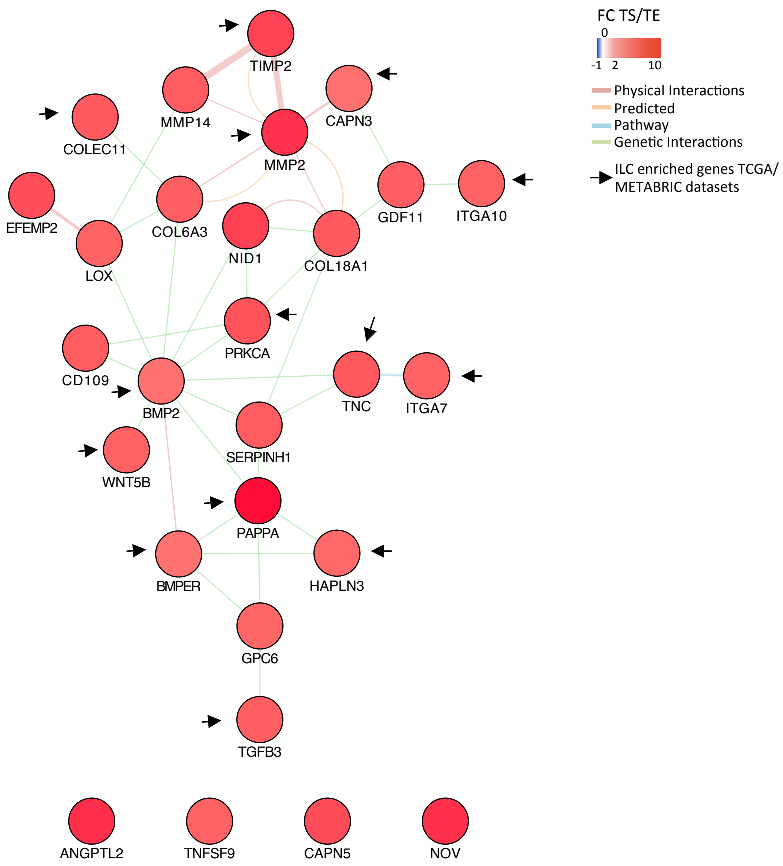


**Supplementary Figure 5: GeneMANIA analysis of CAF associated genes.** Interaction map of the 28 genes that were expressed in the primary CAF dataset was generated using GeneMANIA in Cytoscape. Colour represents the fold change (FC) expression in the TS compared to TE. Blue = FC TS/TE -1 – 0; White = FC TS/TE 0; Pink = FC TS/TE 1 – 2; Red = FC TS/TE 2 – 10. Network connectors represent physical interactions (pink), predicted (orange), pathway (blue) or genetic interactions (green). Arrows point to the 14 genes whose expression was significantly increased in ILC compared to IDC in TCGA and METABRIC datasets.

Supplementary Figure 6

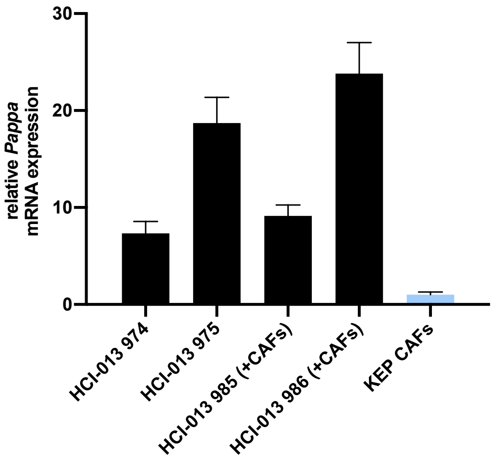


**Supplementary Figure 6: Expression of *Pappa* in ILC patient derived xenograft HCI-013**. qPCR of mouse Pappa in 4 HCI-013 patient-derived xenografts grown in mice in the presence or absence of human CAFs. Pappa expression detected in HCI-013 xenografts is of mouse origin, indicating that it is expressed by mouse fibroblasts present in the tumor stroma. Human PAPPA was not detected in any of the tumors. Pappa in mouse CAFs from the KEP model shown for comparison. Results are shown as the relative expression in KEP-CAFs.

Supplementary Figure 7

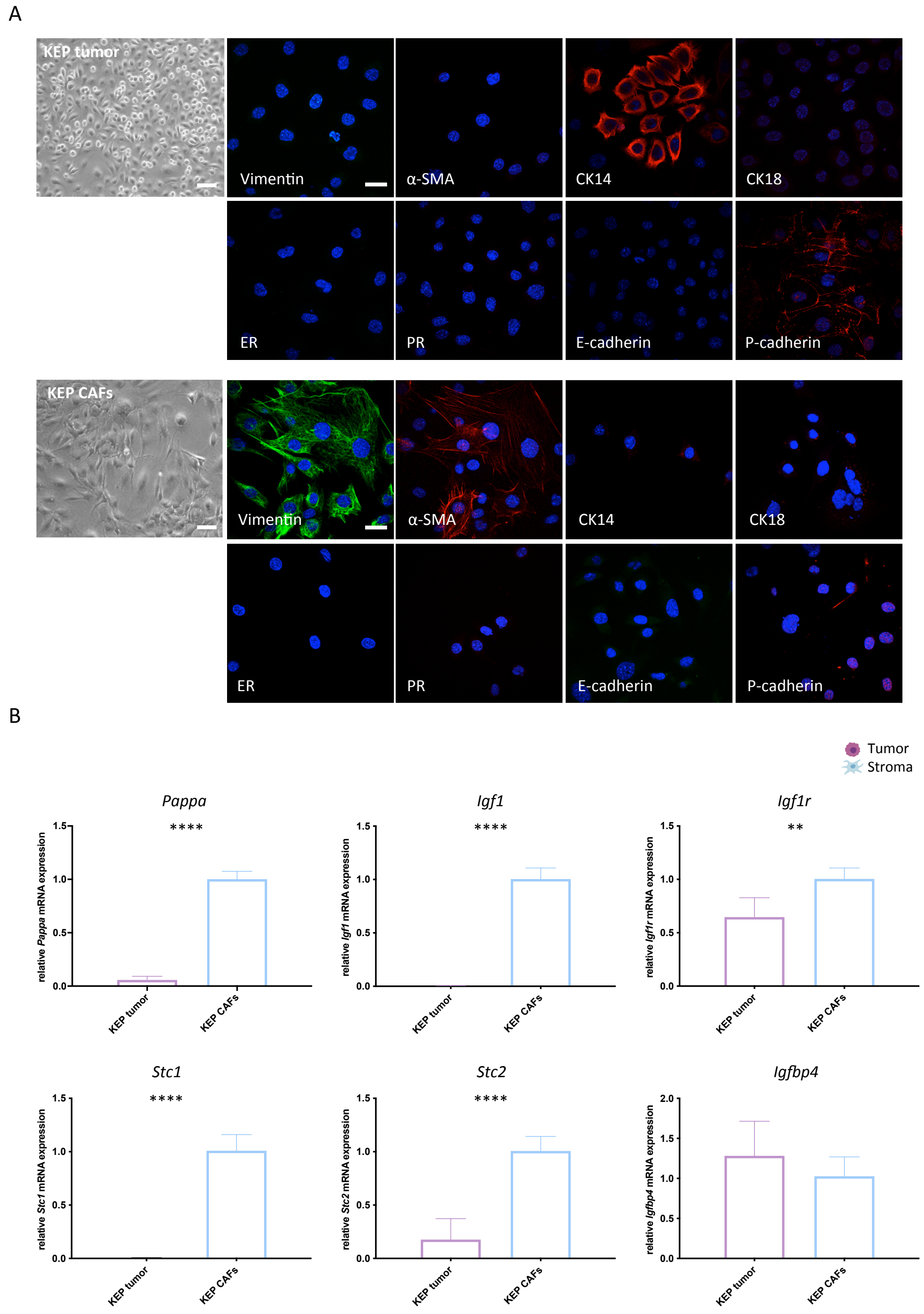


**Supplementary Figure 7: Characterization of ILC KEP mouse model.** A) Phase contrast and confocal images of KEP derived epithelial tumor cells (KEP-tumor) and KEP derived fibroblasts (KEP-CAFs). Cells were stained for vimentin, a-SMA, cytokeratin 14 (CK14), cytokeratin 18 (CK18), estrogen receptor (ER), progesterone receptor (PR), E-cadherin and P-cadherin. Phase contrast scale bars = 100 µm. Confocal scale bars = 30 µm. B) qPCR of Pappa and related genes in KEP-derived epithelial cells and CAFs. Results are shown as the relative expression in KEP-CAFs. Graphs were generated in Graph Pad. ** $p \leq 0.01$; **** $p \leq 0.0001$, t-test comparing expression in KEP-CAFs to KEP tumor.

Supplementary Figure 8

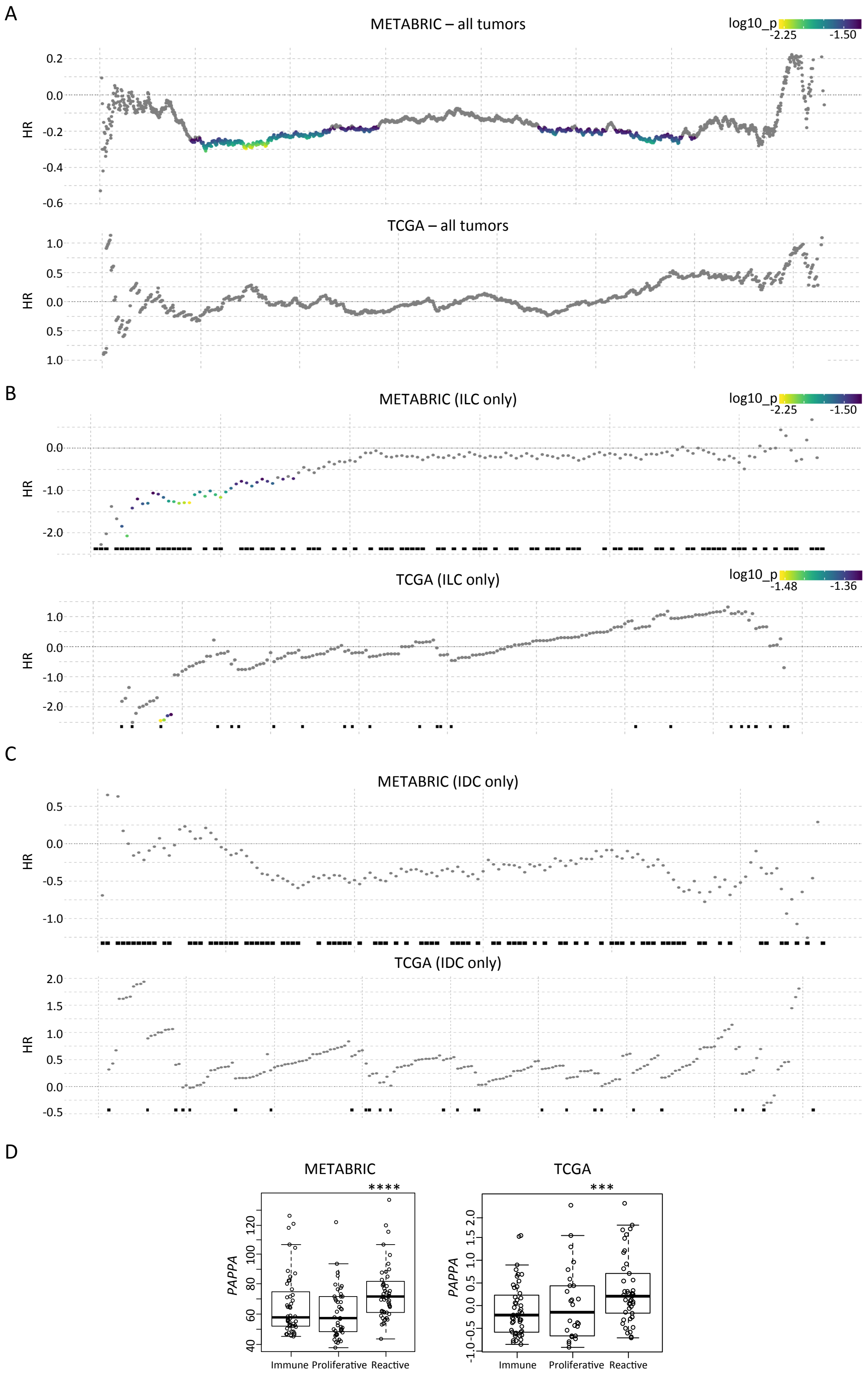


**Supplementary Figure 8: Comprehensive survival analysis of *PAPPA* highlights low expression in ILC, but not IDC is associated with poor outcome.** A) Hazard ratios of all possible (n-1) cut-points to split the 1904 breast tumors of the METABRIC dataset and 1098 breast cancers of the TCGA dataset into two groups ranked left-right by PAPPA expression. Cox proportional hazards p-values less than 0.05 are highlighted by yellow-blue colours. B) survivALL plots for PAPPA expression in ILC tumors from the METABRIC and TCGA studies. C) Representative survivALL plots for 142 and 206 randomly sampled ER+ IDCs from METABRIC and TCGA to compare with the same number of ILCs shown in Supplementary Figure 8B. survivALL plots performed to examine all possible points of separation to split cohorts into low and high groups. D) PAPPA is expressed at significantly higher levels in tumors assigned to the "reactive" ILC subtype (as defined by Ciriello G., et al 2015) in both datasets.

#### References


1. Johnson WE, Li C, Rabinovic A. Adjusting batch effects in microarray expression data using empirical Bayes methods. Biostatistics 2007;8(1):118-27.
2. Oh EY, Christensen SM, Ghanta S*, et al.* Extensive rewiring of epithelial-stromal co-expression networks in breast cancer. Genome Biol 2015;16:128.
3. Finak G, Sadekova S, Pepin F*, et al.* Gene expression signatures of morphologically normal breast tissue identify basal-like tumors. Breast Cancer Res 2006;8(5):R58.
4. Gyrup C, Christiansen M, Oxvig C. Quantification of proteolytically active pregnancy-associated plasma protein-A with an assay based on quenched fluorescence. Clin Chem 2007;53(5):947-54.
5. Gyrup C, Oxvig C. Quantitative analysis of insulin-like growth factor-modulated proteolysis of insulin-like growth factor binding protein-4 and -5 by pregnancy-associated plasma protein-A. Biochemistry 2007;46(7):1972-80.
